# An oncolytic adenovirus armed with anticancer prodrug-activating enzyme offers enhanced tumor killing and antitumor immunity

**DOI:** 10.64898/2026.05.26.727470

**Authors:** Mengshi Sun, Suhua Guan, Chenchen Yang, Hongyang Zhang, Dexiang Xu, Hengchun Li, Pingchao Li, Chunhua Wang, Jin Li, An Hong, Linbing Qu, Ling Chen

## Abstract

Oncolytic viruses are most commonly administered via intratumoral injection; however, their clinical efficacy in achieving tumor eradication remains limited by several challenges, including insufficient penetration into all tumor cells and the inability to elicit robust systemic antitumor immune responses capable of eliminating metastatic microtumors. Here, we report an oncolytic adenovirus, OAd-2B6, with an engineered adenoviral E1 region for tumor selectivity and carrying the prodrug-activating enzyme cytochrome P450 2B6 (CYP2B6) to activate the anticancer prodrug cyclophosphamide (Cytoxan, CTX). OAd-2B6 alone induced dose-dependent tumor cell killing across multiple human tumor cell lines and exhibited strong synergistic antitumor effects when combined with CTX. Importantly, OAd-2B6-mediated local activation of CTX resulted in a potent bystander killing effect that eliminated tumor cells not directly infected by the virus. In a H1299 lung cancer xenograft nude mouse model, intratumoral injection of OAd-2B6 combined with CTX significantly inhibited tumor growth and even achieved complete tumor regression, with markedly superior efficacy compared with monotherapy. In immunocompetent mice bearing 4T1 breast cancer xenografts, OAd-2B6 alone inhibited tumor growth and was accompanied by upregulation of IFN-γ and GzmB expression in the tumor-infiltrated T cells. CTX combination therapy further enhances this anti-tumor immune response, promoting the activation of T cells to suppress non-injected tumors at a distal site. Collectively, this study demonstrates that OAd-2B6 exerts potent antitumor effects through multiple mechanisms, including direct oncolysis, intratumoral prodrug activation leading to bystander killing, and enhancement of systemic antitumor immunity. These findings provide a promising strategy for improving the therapeutic efficacy of oncolytic therapy.

## Introduction

With intrinsic tumor-specific lysis capacity and immunostimulatory properties, oncolytic viruses have emerged as promising therapeutic agents for immunotherapy of solid tumors[1]. Since the phenomenon of virus-mediated tumor regression was confirmed in the early 20th century[2], a variety of viral vectors have entered clinical research and development, and four oncolytic virus products have been officially approved for clinical use worldwide[3–5]. Nevertheless, the clinical application of current oncolytic viruses in solid tumors remains limited. The dense extracellular matrix and immunosuppressive tumor microenvironment collectively form physical and immune barriers[6]. These barriers not only restrict the deep penetration and diffusion of viruses within the tumor parenchyma, but also hinder intratumoral viral replication and anti-tumor immune activation, ultimately resulting in a low clinical response rate to single-agent oncolytic virus treatment and limiting its efficacy against refractory solid tumors[7].

Among various oncolytic viral vectors, adenoviruses are considered one of the most mature vectors for clinical translation due to their easy modification, large packaging capacity, and non-integrating characteristics into the host genome[8]. Clinical data have demonstrated that engineered oncolytic adenoviruses exert definite therapeutic effects on multiple tumor types [9–11]. For instance, H101 has been shown to markedly improve the chemotherapy response rate in solid tumors [12,13], and CG0070 exhibits favorable safety and efficacy in non-muscle-invasive bladder cancer [14]. However, multiple adenovirus candidates have encountered obstacles in clinical research. As exemplified by Onyx-015, insufficient in vivo replication efficiency, weak tumor selectivity, and poor anti-tumor activity impeded its progression to late-stage clinical trials [15]. Collectively, tumor matrix densification and immune cold phenotype remain the core bottlenecks restricting the efficacy of oncolytic adenoviruses against solid tumors. Therefore, the development of rational combination therapeutic strategies to remodel the tumor microenvironment is critical to break through the limitations of oncolytic virus therapy[16].

Cyclophosphamide (CTX), a commonly used alkylating agent, exhibits dose-dependent dual pharmacological effects: low-dose CTX alleviates tumor-induced immunosuppression, whereas high-dose CTX induces DNA alkylation damage in tumor cells, inhibiting tumor proliferation and producing a bystander-killing effect to eliminate adjacent tumor cells [17,18]. Such dual properties endow CTX with inherent synergistic potential when combined with oncolytic viruses. CTX is primarily metabolized and activated in the liver by the cytochrome P450 enzyme system. CYP2B6 serves as a pivotal subtype regulating CTX activation in humans, which catalyzes the generation of active metabolite phosphoramide mustard to exert anti- tumor effects[16,19]. Nevertheless, systemic administration leads to non-specific hepatic activation of CTX, resulting in poor tumor targeting and dose-limiting toxicities such as myelosuppression and nephrotoxicity. To mitigate systemic adverse effects and achieve tumor-localized targeted activation of CTX, our previous studies have confirmed that specific overexpression of CYP450 enzymes in tumor cells enhances the chemosensitivity of tumor cells to CTX in a dose-dependent manner and realizes in-situ drug activation, validating the feasibility of enzyme-mediated prodrug activation[20]. However, viral vectors currently used for enzyme delivery are mostly replication-deficient viruses, which exhibit weak tumor targeting and short intratumoral retention time, thus failing to effectively treat dense solid tumors and limiting their clinical translational value [21]. In summary, CTX is accompanied by non-specific systemic activation toxicity, and conventional gene delivery vectors are not suitable for the combined therapeutic system of oncolytic viruses.

To overcome the above technical bottlenecks, a novel replication-competent oncolytic adenovirus OAd-2B6 stably expressing human-derived CYP2B6 was constructed based on the clinically isolated recombinant adenovirus backbone HAdV- C104[22]. This engineered vector possesses the dual advantages of oncolytic virus and enzyme-mediated prodrug therapy. Genetically modified adenovirus harbors tumor- selective replication capability to lyse malignant tumor cells. Meanwhile, exogenous CYP2B6 mediates intratumoral targeted drug activation, thereby enhancing chemotherapeutic killing efficacy and reducing systemic toxic side effects. Furthermore, this combined treatment synergistically strengthens the systemic anti-tumor immune response and expands the range of intratumoral killing via the bystander effect. In this study, lung solid tumor models were established to explore the anti-tumor efficacy and immune regulatory mechanism of OAd-2B6 combined with CTX. In conclusion, this research develops a combined therapeutic strategy that integrates intratumoral targeted activation and immune synergy, aiming to provide a theoretical basis for targeted combination therapy for solid tumors and to lay a research foundation for the clinical transformation and application of oncolytic viruses.

## Results

### Design and Construction of the CYP2B6-Armed Conditionally Replicating Oncolytic Adenovirus OAd-2B6

We designed and constructed an oncolytic adenovirus Ad104-2B6 (OAd-2B6) based on the clinically isolated human adenovirus type C strain HAdV-C104, which contains the penton and hexon genes of human adenovirus type 1 and the fiber gene of human adenovirus type 2[23]. In order to enhance the specificity and selectivity for tumors as well as the anti-tumor activity, the E1-CR2 region related to the proliferation of normal host cells was deleted[24,25]. Furthermore, previous studies have revealed that knockout of the anti-apoptotic E1B-19K gene capable of binding Beclin-1 and restraining apoptosis can further strengthen p53-induced apoptotic effects in tumor cells[26–28]. The immunosuppressive E3 region was also deleted and replaced with the CYP2B6 gene, which encodes a prodrug-activating enzyme capable of converting the anticancer prodrug CTX into its active cytotoxic metabolites (Fig. 1a). This design not only aimed to enable selective replication of oncolytic adenovirus in tumor cells but also enable the expression of prodrug activation enzyme in the tumor cells, rendering tumor cells to activate CTX within the tumor microenvironment.

**Fig 1:**
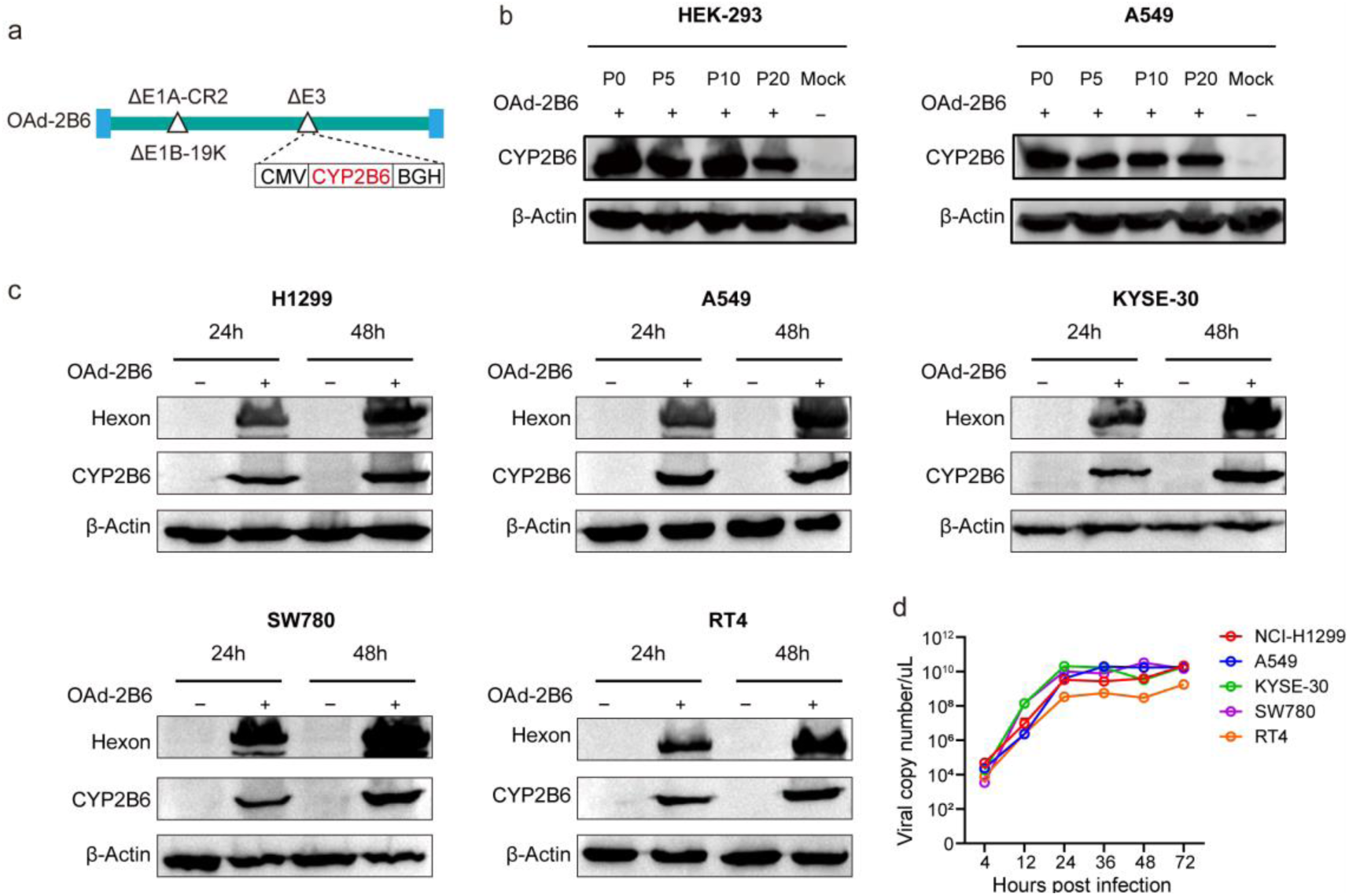
Construction and basic biological features of OAd-2B6. **a.** Schematic structure of the OAd-2B6 vector. The E1B-19K sequence was deleted to strengthen the oncolytic capacity, and the human CYP2B6 gene was inserted into the E3 region to activate the prodrug CTX. **b.** HEK293 cells were used for continuous viral passage. Viral supernatants at passages 5, 10, and 20 were used to infect HEK293 and A549 cells. CYP2B6 protein expression was detected by Western blot. **c.** Tumor cell lines were infected with OAd-2B6 at an MOI of 3. Viral hexon and CYP2B6 protein levels were analyzed by Western blot at 24 h and 48 h post-infection. **d.** Viral replication was evaluated by quantifying adenoviral E1 gene copy numbers using qPCR in H1299, A549, KYSE-30, SW780 and RT4 cells infected with OAd-2B6 at an MOI of 3 at indicated time points post-infection (4-72 h).

To evaluate the genetic stability of the newly constructed oncolytic adenovirus, OAd-2B6 was serially passaged for 20 generations. Viral stocks collected from passages 5, 10, and 20 were used to assess exogenous CYP2B6 expression in HEK293 and A549 cells. The results demonstrated that OAd-2B6 maintained stable CYP2B6 expression throughout at least 20 serial passages without detectable gene loss, indicating favorable genomic stability of the recombinant virus (Fig. 1b).

To further investigate the infectivity and exogenous CYP2B6 expression profile of OAd-2B6 in tumor cells, multiple human cancer cell lines, including H1299, A549, KYSE-30, SW780, and RT4, were infected with OAd-2B6 at 3 MOI. Western blot analysis revealed detectable expression of both adenoviral hexon and CYP2B6 proteins as early as 24 h post-infection, with expression levels further increased at 48 h. These findings confirmed that OAd-2B6 efficiently infected a broad spectrum of tumor cells and underwent active intracellular replication, demonstrating broad tumor tropism and replication capability across multiple cancer cell types (Fig. 1c). To evaluate the replication capability of OAd-2B6 in human tumor cells, progeny viral production was quantified at different time points post-infection. The results showed that viral copy numbers increased markedly at 24 h post-infection and persisted until 72 h (Fig. 1d), indicating that OAd-2B6 efficiently replicated in multiple human tumor cell lines.

### OAd-2B6 Efficiently Eliminates Human Tumor Cells, and CTX Enhances Its Cytotoxicity Through a Bystander Effect

Next, we investigated the tumor selectivity and cytotoxic effects of OAd-2B6 (Fig. 2a). Human normal bronchial epithelial BEAS-2B cells (Fig. 2b) and human tumor cell lines, including NCI-H1299, A549, SW780, and RT4 (Fig. 2c), were infected with OAd-2B6 at different multiplicities of infection (MOIs). BEAS-2B cells exhibited minimal sensitivity to OAd-2B6, showing no obvious cytotoxicity at 1 MOI, while maintaining approximately 80% viability even after infection at 100 MOI (Fig. 2b), indicating limited toxicity toward normal cells. In contrast, the viability of NCI-H1299, A549, SW780, and RT4 tumor cells decreased significantly in an MOI-dependent manner following OAd-2B6 infection (Fig. 2d). To explore whether OAd-2B6- mediated CYP2B6 expression, followed by activation of the anticancer prodrug CTX, could potentiate tumor killing efficacy. In lung cancer NCI-H1299 cells, treatment with OAd-2B6 alone at an MOI of 0.2 exerted only mild cytotoxicity, with a cell viability of 92%. Combined treatment with 0.1, 0.3 and 0.5 mM CTX reduced cell viability to 69.6%, 15.5% and 13.4%, respectively, resulting in markedly enhanced cytotoxic effects (Fig. 2d). For lung cancer A549 cells, single treatment with OAd-2B6 at an MOI of 3 led to a cell viability of 79%, while combined administration with 0.1, 0.3 and 0.5 mM CTX decreased cell viability to 48.4%, 41.2% and 33.6% in sequence (Fig. 2d). In bladder cancer SW780 cells, OAd-2B6 monotherapy at an MOI of 0.5 yielded a cell viability of 73.9%, and the viabilities after combined treatment with gradient concentrations of CTX were 64.3%, 32.5% and 3.97% (data to be supplemented). Bladder cancer RT4 cells showed a cell viability of 90.9% after single OAd-2B6 treatment at an MOI of 0.5 (data to be confirmed), and the viabilities dropped to 50.4%, 22.8% and 17.1% when combined with 0.1, 0.3 and 0.5 mM CTX. Compared with the OAd-2B6 monotherapy group, the combined treatment group exhibited remarkably improved tumor suppressive effects, and the anti-tumor activity presented a CTX concentration-dependent manner (Fig. 2d). These results demonstrated that the combination of OAd-2B6 and CTX exerted synergistic cytotoxicity and achieved dual anti-tumor effects.

**Fig 2:**
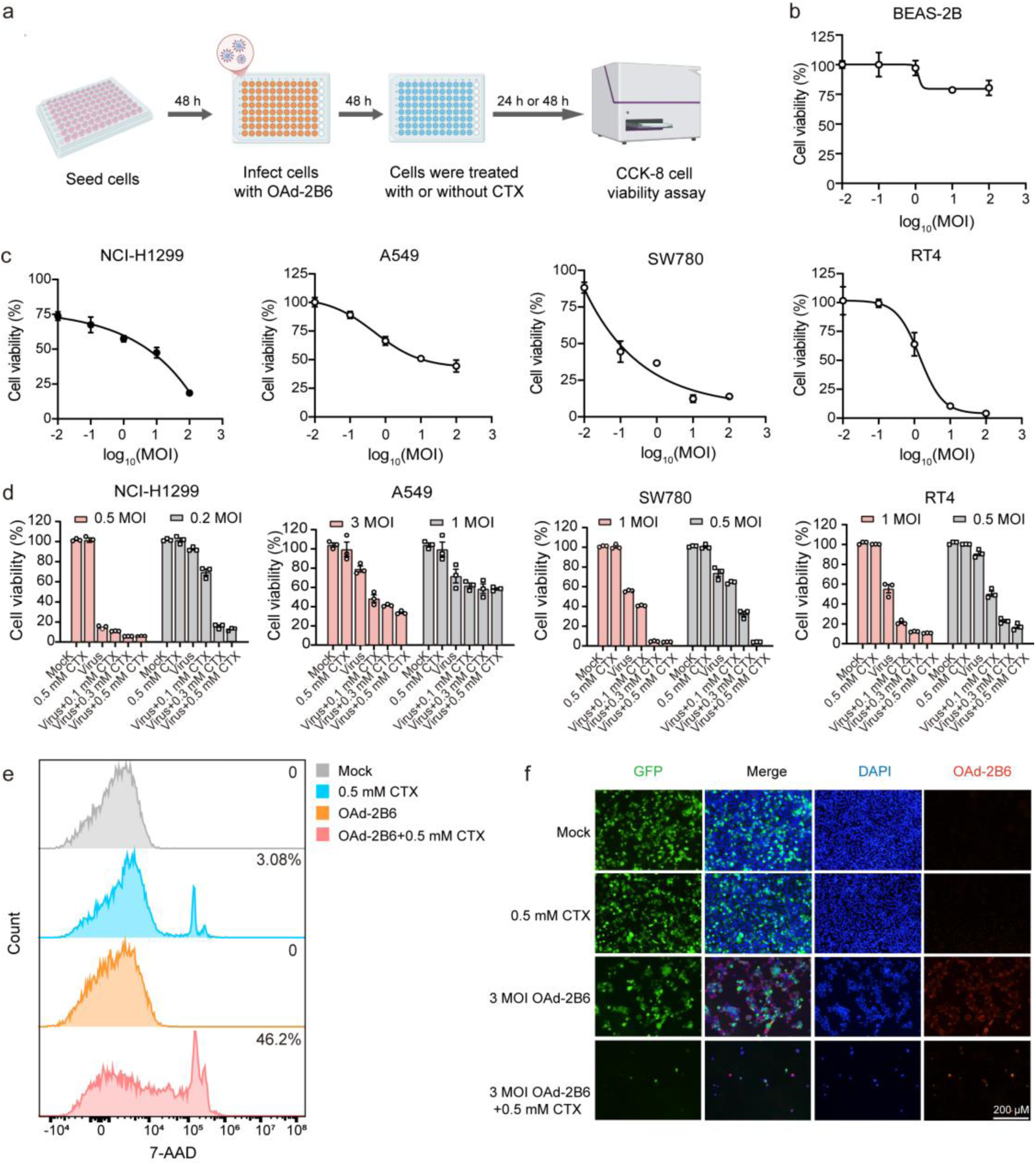
Cyclophosphamide potentiates the inhibitory effect of OAd-2B6 on multiple tumor cell lines via bystander killing effects. **a.** Schematic diagram of the in vitro cytotoxicity experiment of OAd-2B6 monotherapy combined with CTX. **b-c.** BEAS-2B normal epithelial cells (b) and four tumor cell lines (c) were infected with OAd-2B6 at different MOIs. After 72 h of treatment, cell viability was measured using the CCK-8 assay. **d.** Tumor cells were infected with OAd-2B6 at different MOIs and subsequently treated with CTX at gradient concentrations. The changes in cell viability were evaluated by the CCK-8 assay. **e.** H199-GFP cells were incubated with the supernatant derived from OAd-2B6-transfected and CTX-treated H1299 cells. The proportion of PI-positive cells was analyzed by flow cytometry. **f.** Immunofluorescence staining was performed to detect adenoviral Hexon in the co-culture system with or without OAd-2B6 infection and CTX treatment. Fluorescence images were captured using a fluorescence microscope.

### Activation of anticancer prodrug CTX leads to strong bystander killing of tumor cells without OAd-2B6 infection

The inability of oncolytic viruses to penetrate and infect all tumor cells within solid tumors remains a major bottleneck limiting their clinical application. Accumulated preclinical studies have demonstrated that activated cyclophosphamide metabolites can exert potent bystander cytotoxic effects on neighboring tumor cells lacking P450 enzyme expression. To verify whether the combination of OAd-2B6 and CTX mediates the bystander effect, conditioned-medium assays were performed to assess apoptosis in H1299-GFP cells. Compared with the Mock and OAd-2B6 monotherapy groups, only 53.8% of cells remained viable following combined OAd- 2B6 and CTX treatment (Fig. 2e). Furthermore, when OAd-2B6-infected H1299 cells were co-cultured with uninfected H1299-GFP cells, treatment with 0.5 mM CTX markedly reduced the number of H1299-GFP cells compared with the 1 MOI OAd-2B6 monotherapy group, regardless of whether direct infection occurred (Fig. 2f). These results indicate that activated CTX metabolites generated by OAd-2B6-mediated CYP2B6 expression can induce significant bystander killing of neighboring tumor cells.

### Intratumoral Administration of OAd-2B6 Suppresses or Eliminates Xenograft Tumors in Nude Mice

To investigate the *in vivo* antitumor efficacy of OAd-2B6, a human lung cancer H1299 subcutaneous xenograft model was established in nude mice. Intratumoral injections of OAd-2B6 were administered at doses of 3 × 10⁹ vp or 3 × 10¹⁰ vp, and tumor growth was monitored for 3 weeks (Fig. 3a). Tumor volume analysis demonstrated that both low-dose (3 × 10^9^ vp) and high-dose (3 × 10¹⁰ vp) OAd-2B6 treatment significantly inhibited tumor growth compared with the PBS group (Fig. 3b). Furthermore, combined treatment of OAd-2B6 and CTX markedly decreased tumor weight compared with the PBS group (Fig. 3c). Survival analysis further showed that, compared with the PBS group exhibiting a survival rate of 30%, treatment with low-dose OAd-2B6 increased survival to 90%. In contrast, all mice in the high-dose OAd- 2B6 group survived throughout the observation period (Fig. 3d), indicating that OAd- 2B6 monotherapy effectively prolonged survival by suppressing tumor progression.

**Fig 3:**
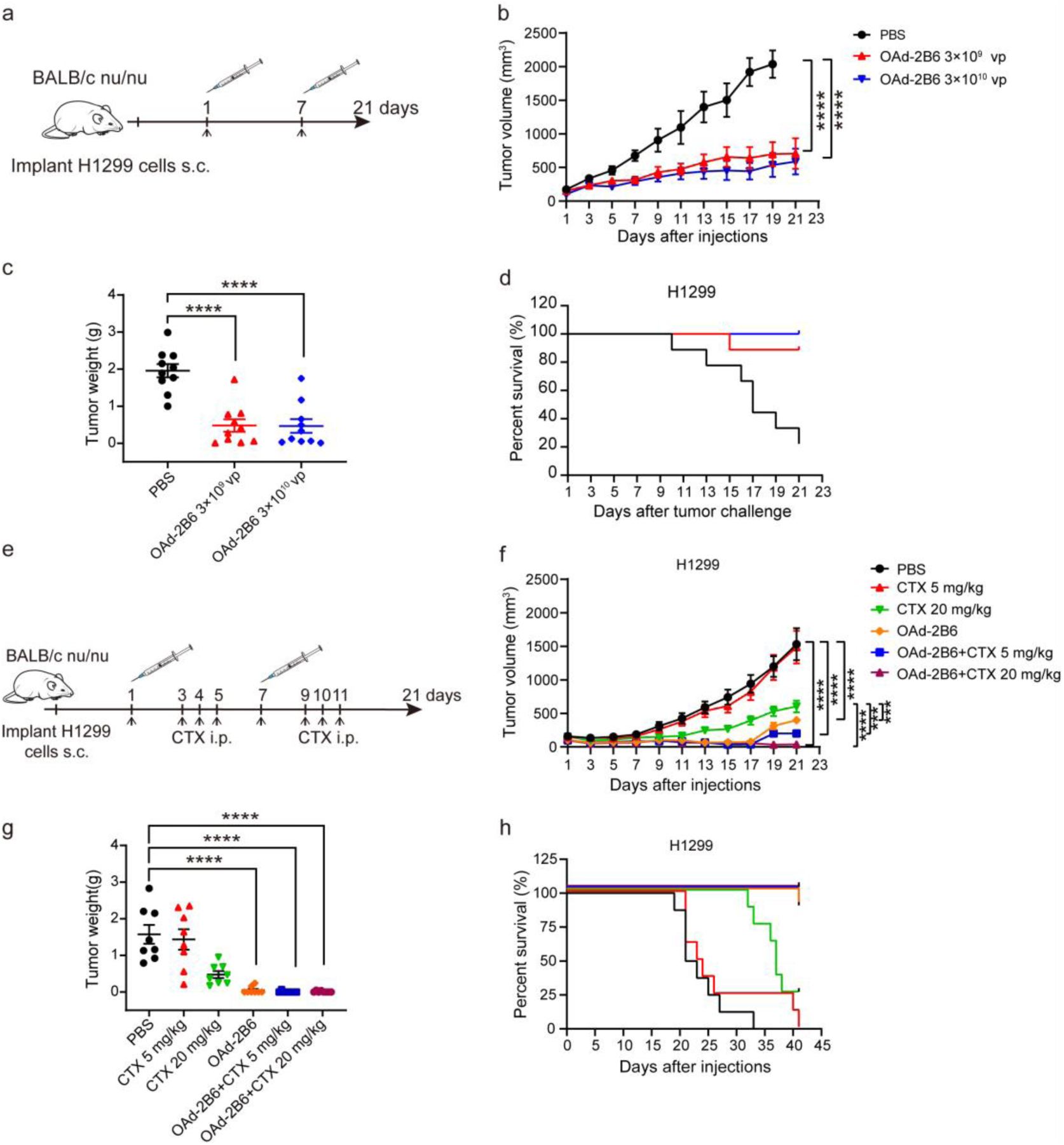
Intratumoral administration of OAd-2B6 inhibits xenograft tumor growth and prolongs the survival time of nude mice. **a.** Schematic diagram of the single-treatment regimen of OAd-2B6 in the H1299 xenograft model. Mice were intratumorally injected with OAd-2B6 at doses of 3×10⁹ vp and 3×10¹⁰ vp per mouse, with an identical injection volume of 50 μL per animal on days 1 and 7 after tumor formation. **b.** Average tumor growth curves of H1299 tumor-bearing mice (n=10). **c.** Tumor weight of each group was quantified on day 21 after treatment. **d.** Survival curves of tumor-bearing mice. **e.** Schematic diagram of the combined treatment regimen of OAd-2B6 and CTX. 48 hours after viral injection, CTX was administered intraperitoneally for 3 consecutive days, and the treatment cycle was repeated after 7 days (n=8). **f.** Tumor growth curves of mice in monotherapy and combination groups (n=8). **g.** Tumor weight of mice from different treatment groups. **h.** Survival rate of mice in each group (n=8). Data are presented as mean ± SEM. *P < 0.05, **P < 0.01, ***P < 0.001, ****P < 0.0001.

To further validate the synergistic antitumor effects of OAd-2B6 combined with cyclophosphamide in vivo (Fig. 3e), tumor growth was monitored following combination treatment. Compared with the PBS and 20 mg/kg CTX groups, OAd-2B6 monotherapy significantly reduced tumor volume, while the combination of OAd-2B6 and CTX produced even more pronounced tumor suppression, with complete tumor regression observed in some mice (Fig. 3f). Consistently, tumor weights at day 21 were significantly lower in the OAd-2B6 monotherapy group as well as in the OAd-2B6 plus CTX combination groups (Fig. 3g).

To further clarify the synergistic therapeutic effect of CTX on OAd-2B6 in vivo, survival monitoring was extended to 41 days. By day 33, all mice in the PBS group had died, with a median survival time of only 22 days, indicating the aggressive progression of untreated H1299 xenografts. Treatment with 5 mg/kg CTX modestly prolonged survival to 41 days and increased the median survival time to 23.5 days. At the same time point, survival rates in the 20 mg/kg CTX group and OAd-2B6 monotherapy group were 25% and 87.5%, respectively. Although high-dose CTX extended the median survival time to 37 days, OAd-2B6 monotherapy improved overall survival by approximately 3.5-fold compared with CTX alone (Fig. 3h), demonstrating superior therapeutic efficacy in improving survival outcomes.

### Intratumoral Administration of OAd-2B6 Combined with CTX Suppresses Tumor Growth and Enhances T Cell Infiltration in Immunocompetent Mice

To evaluate the therapeutic efficacy of the oncolytic virotherapy in a poorly immunogenic tumor model, a highly metastatic and immunosuppressive 4T1 breast cancer model was employed to assess the antitumor activity of OAd-2B6 in immunocompetent mice (Fig. 4a). The results showed that, compared with the PBS group, OAd-2B6 monotherapy significantly reduced tumor volume and demonstrated superior efficacy compared with cyclophosphamide (CTX, 20 mg/kg) alone. Notably, combined treatment with OAd-2B6 and CTX resulted in a markedly greater tumor growth inhibition than OAd-2B6 monotherapy, indicating a clear synergistic antitumor effect and suggesting that CTX enhances the therapeutic efficacy of OAd-2B6 (Fig. 4b). In addition, both OAd-2B6 monotherapy and the combination regimen significantly delayed tumor progression and reduced tumor weight (Fig. 4c), indicating that this combination strategy exerts favorable in vivo antitumor efficacy.

**Fig 4:**
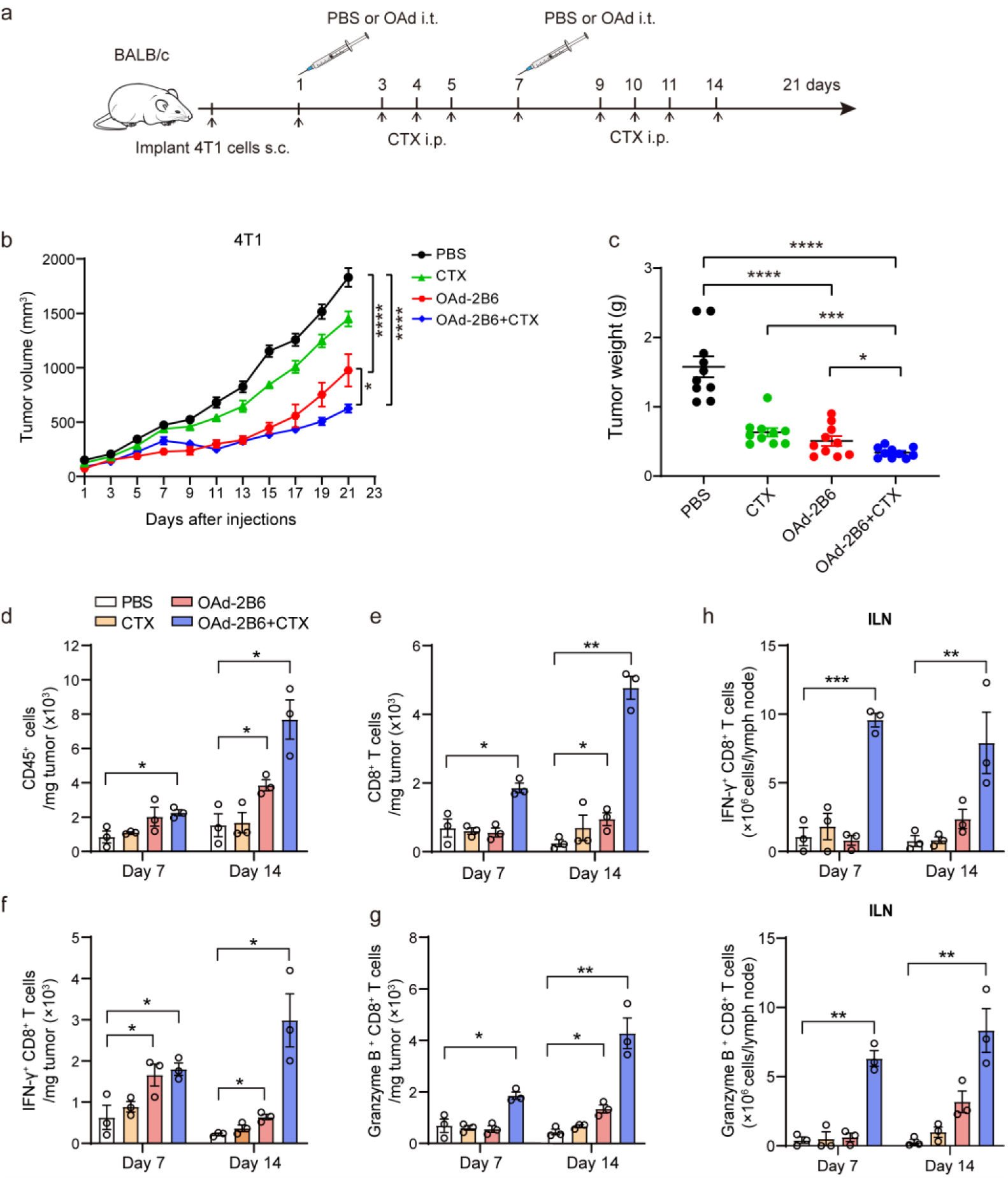
Combined treatment with OAd-2B6 and cyclophosphamide inhibits tumor growth by elevating intratumoral T lymphocyte infiltration. **a.** Schematic illustration of the treatment schedule in the 4T1 tumor-bearing mouse model. Mice were intratumorally injected with 1×10^11^ vp OAd-2B6 (100 μL per mouse) on days 1 and 7 post-tumor formation. Subsequently, mice received intraperitoneal injection of 20 mg/kg CTX (100 μL per mouse) on days 3, 4, 5, 9, 10, and 11 after treatment initiation (n=10). **b.** Tumor growth curves of 4T1 breast cancer-bearing mice (n=10). **c.** Tumor weight of the treated tumors in each group collected at the experimental endpoint. **d-g.** Female BALB/c mice aged 5-6 weeks were inoculated with 4T1 cells. Mice were treated with intratumoral injection of OAd-2B6 alone or combined with cyclophosphamide. The mice were euthanized on day 7 and day 14 post-treatment. Flow cytometry was used to detect the number and function of immune cells in the untreated tumor microenvironment, including absolute counts of tumor-infiltrating CD45⁺ (d) and CD8⁺ T cells (e), as well as the proportions of IFN-γ⁺ (f) and Granzyme B⁺ (g) cells among CD8⁺ T cells (n=3). Data are presented as mean ± SEM. *P < 0.05, **P < 0.01, ***P < 0.001, ****P < 0.0001. **h.** Expression levels of IFN-γ⁺ and Granzyme B⁺ in CD8⁺ T cells of axillary lymph nodes.

Beyond direct oncolysis, oncolytic viruses exert antitumor effects primarily by activating local immune responses and remodeling the tumor immune microenvironment. To further characterize the immunomodulatory effects of OAd-2B6 in the 4T1 model, tumor-infiltrating immune cells were analyzed by flow cytometry on days 7 and 14 after treatment. On day 7, the combination group exhibited a significant increase in the absolute numbers of CD45⁺ immune cells and CD8⁺ T cells within the tumor microenvironment, and this increase continued through day 14. In contrast, significant increases in CD45⁺ and CD8⁺ T cells in the OAd-2B6 monotherapy group were only observed at day 14 (Fig. 4d, e). These findings indicate that combination therapy induces a more rapid and robust immune activation, whereas OAd-2B6 alone may require a longer time to initiate local immune responses.

Consistently, the proportion of effector CD8⁺ T cells was significantly elevated in the combination group as early as day 7. By day 14, both OAd-2B6 monotherapy and combination therapy significantly increased the numbers of IFN-γ⁺ CD8⁺ T cells and Granzyme B⁺ CD8⁺ T cells, with the combination group showing a markedly stronger effect than monotherapy (Fig. 4f, g). These results indicate that OAd-2B6 effectively activates tumor-specific CD8⁺ T cells and that co-administration with CTX further enhances cytotoxic T cell responses and antitumor immunity.

As tumor-draining lymph nodes are critical sites for initiating systemic immune responses, we further analyzed functional CD8⁺ T cells in the inguinal lymph nodes. Both OAd-2B6 monotherapy and combination treatment significantly increased the absolute numbers of IFN-γ⁺ CD8⁺ T cells and Granzyme B⁺ CD8⁺ T cells in the draining lymph nodes (Fig. 4h). The kinetics of CD8⁺ T cell activation in lymph nodes closely paralleled the observed tumor regression, indicating that immune activation in lymph nodes is crucial for initiating potent local anti-tumor immune responses.

### OAd-2B6 Combined with CTX Induces Systemic Antitumor Immune Responses and Suppresses Distant Tumor Growth in Immunocompetent Mice

Tumor-specific antitumor immune responses are critical for long-term tumor suppression and distant tumor inhibition. Immune activation in draining lymph nodes effectively initiates systemic antitumor immunity and mediates abscopal antitumor effects[29,30]. Therefore, this study further investigated whether OAd-2B6 combined with CTX could activate systemic specific immunity and exert distant tumor-inhibitory effects. To determine whether OAd-2B6 monotherapy or its combination with cyclophosphamide (CTX) could induce tumor antigen-specific T cell responses in vivo, an IFN-γ enzyme-linked immunospot (ELISPOT) assay was performed to measure IFN-γ secretion by splenocytes stimulated with tumor cell lysates. The results showed that both the OAd-2B6 monotherapy group and the OAd-2B6+CTX combination group exhibited significantly higher numbers of IFN-γ-positive spot-forming cells compared with the PBS and CTX monotherapy groups. Moreover, the combination treatment further increased the number of IFN-γ spots compared with OAd-2B6 alone (Fig. 5a, b).

**Fig 5:**
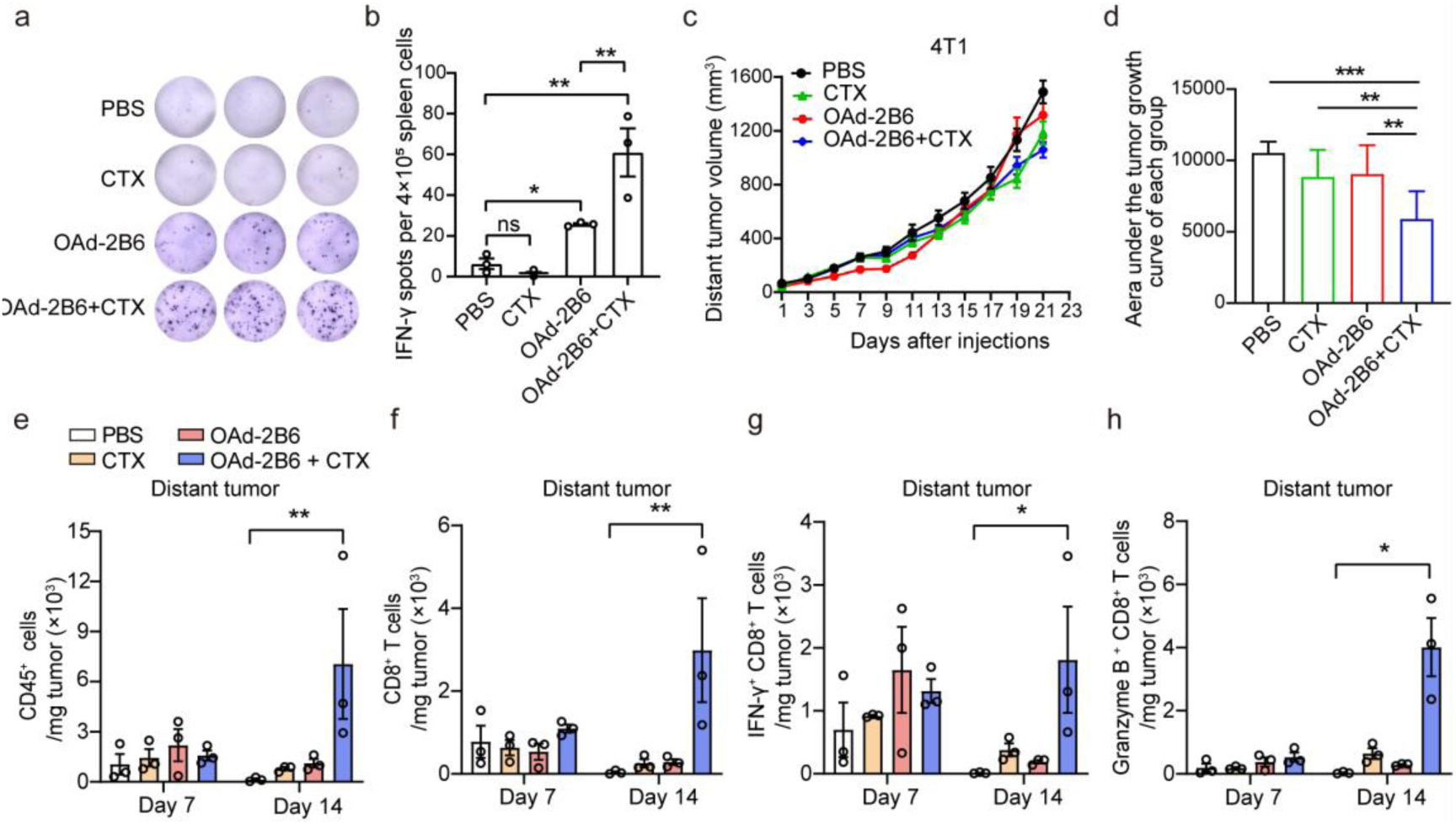
Intratumor injection of OAd-2B6 with CTX treatment in immunocompetent mice elicits antitumor immunity against distal tumors. **a.** After stimulation with 4T1 tumor lysate, 4×10^5^ splenocytes were used for IFN-γ ELISPOT assay to observe spot formation (n=3). **b.** The number of ELISPOT spots was quantified to evaluate the IFN-γ secretion level of splenocytes. **c.** Growth curves of distant untreated tumors in 4T1 tumor-bearing mice (n=10). **d.** Statistical analysis of area under the curve for the volume of distant untreated tumors. **e-g.** Flow cytometry was applied to quantify the proportions of CD45^+^ immune cells (e), CD8^+^ T cells (f), IFN-γ^+^ CD8^+^ T cells (g), and Granzyme B^+^ CD8^+^ T cells (h) in untreated tumors. (n=3). Data are presented as mean ± SEM. *P < 0.05, **P < 0.01, ***P < 0.001, ****P < 0.0001.

We next evaluated the antitumor effect of unilateral intratumoral administration on contralateral (distant) tumors. At day 21 post-treatment, both OAd-2B6 and CTX monotherapy groups inhibited the growth of distant tumors, whereas the combination therapy exhibited the most pronounced tumor-suppressive effect (Fig. 5c). Meanwhile, AUC analysis was performed on the volume of distant untreated tumors in each group. The results revealed that the area under the tumor growth curve was markedly lower in the OAd-2B6 combined with CTX group compared with the CTX monotherapy group and OAd-2B6 single-agent group, indicating that the combination treatment could effectively inhibit the growth of distant untreated tumors and exert potent abscopal antitumor effect (Fig. 5d). In addition, on day 14 after treatment, a significant increase in CD45⁺ immune cell infiltration was observed in distant tumors of the combination group, while no significant changes were detected in the OAd-2B6 monotherapy group at either early or late time points (Fig. 5e), suggesting that only the combination regimen effectively activates immune responses in distant tumor sites. Further analysis revealed a marked increase in CD8⁺ T cell infiltration in distant tumors in the combination group, whereas CD8⁺ T cell levels in both the OAd-2B6 and CTX monotherapy groups remained at baseline levels (Fig. 5f). Moreover, combination therapy significantly increased the densities of IFN-γ⁺ CD8⁺ T cells and Granzyme B⁺ CD8⁺ T cells in distant tumors (Fig. 5g, h), indicating that the combined treatment not only promotes infiltration of CD8⁺ T cells into distant tumor sites but also enhances their cytotoxic effector functions.

## Discussion

Oncolytic adenoviruses have emerged as a promising platform for cancer immunotherapy due to their ability to selectively replicate in tumor cells and remodel the tumor immune microenvironment[31,32]. However, their clinical efficacy remains limited by two major barriers: the dense extracellular matrix that restricts intratumoral viral dissemination, and the immunosuppressive tumor microenvironment that impairs effective antitumor immune activation[33]. To address these challenges, we engineered a recombinant oncolytic adenovirus, OAd-2B6, which selectively expresses the cytochrome P450 enzyme CYP2B6 within tumor cells, enabling intratumoral activation of the prodrug cyclophosphamide (CTX) and thereby enhancing local cytotoxicity through a bystander-killing effect.

In vitro, OAd-2B6 demonstrated dose-dependent cytotoxicity across multiple human cancer cell lines and exhibited strong synergistic effects when combined with CTX. In vivo, OAd-2B6 significantly inhibited tumor growth in H1299 xenograft models and achieved complete tumor regression in a subset of treated mice. Notably, compared with previously reported conditionally replicative adenoviruses such as CG0070[34], which require multiple intratumoral administrations to achieve tumor control, OAd-2B6 achieved comparable antitumor efficacy with fewer injections, suggesting potential advantages in therapeutic efficiency.

Prodrug-activating strategies represent an effective approach to enhance oncolytic virotherapy. Classical systems such as herpes simplex virus thymidine kinase/ganciclovir (HSV-TK/GCV) have demonstrated clinical benefit by converting non-toxic prodrugs into cytotoxic metabolites that induce DNA damage and immunogenic cell death[35]. Similarly, cyclophosphamide is a widely used alkylating prodrug whose activation depends on cytochrome P450 enzymes. However, CYP450 expression is generally low in tumor tissues, limiting local CTX activation. In this study, by deleting the E1A-CR2 region to confer tumor-selective replication and inserting CYP2B6 into the E3 locus, OAd-2B6 enables efficient intratumoral conversion of CTX into its active metabolite phosphoramide mustard. This strategy not only enhances local cytotoxicity but also spatially restricts chemotherapy activation to the tumor site, thereby potentially reducing systemic toxicity.

Importantly, OAd-2B6 combined with CTX may help overcome two fundamental barriers of oncolytic virotherapy: physical tumor penetration and immune suppression. The virus-mediated local production of active CTX metabolites enables a potent bystander effect that extends cytotoxicity beyond infected cells, thereby partially compensating for limited viral spread within the tumor mass. Meanwhile, CTX-induced immunomodulation may further reshape the tumor immune microenvironment, facilitating immune cell infiltration and activation. In the immunocompetent 4T1 model, OAd-2B6 alone increased intratumoral CD8⁺ T cell infiltration. At the same time, combination therapy with CTX accelerated and amplified this immune activation, suggesting a synergistic effect in converting “cold” tumors into more inflamed phenotypes.

Despite these promising findings, several limitations should be acknowledged. First, the spatial distribution of viral dissemination and the extent of bystander killing within tumor tissues were not directly quantified, leaving the intratumoral penetration dynamics incompletely characterized. Second, long-term antitumor immune memory and potential protection against tumor rechallenge were not evaluated. Finally, inherent limitations of adenoviral vectors, including pre-existing immunity and rapid clearance in vivo, may restrict therapeutic durability. Future studies incorporating capsid engineering, tumor-targeting modifications, or immune evasion strategies may further improve the in vivo performance of OAd-2B6.

In summary, this study demonstrates that OAd-2B6 in combination with cyclophosphamide exerts potent antitumor effects through coordinated mechanisms, including oncolytic tumor lysis, intratumoral prodrug activation, bystander killing, and immune microenvironment remodeling. This multifunctional design provides a promising strategy to overcome both physical and immunological barriers in solid tumors. It offers a potential platform for next-generation engineered oncolytic adenovirus-based cancer therapy.

## Materials and Methods

### Cell lines

NCI-H1299 and 4T1 cells were cultured in RPMI 1640 medium (Gibco). A549, SW780, 293T and HEK293 cells were maintained in DMEM (Gibco), while RT4 cells were cultured in McCoy’s 5A medium. All media were supplemented with 10% fetal bovine serum (FBS) and 1% penicillin-streptomycin. Cells were incubated at 37°C in a humidified atmosphere containing 5% CO_2_.

### Virus production and purification

The Ad104 virus was originally isolated from clinical samples. For viral amplification, infected cells were harvested when 80-90% of cells showed cytopathic effect (CPE). Cells were collected in sterile 50 mL conical tubes and subjected to three cycles of freeze-thaw lysis. The lysates were centrifuged at 12,000 × g for 10 min at 4°C. The supernatant was carefully collected, and cell debris was discarded. Viral particles were further purified using cesium chloride (CsCl) density gradient ultracentrifugation.

### Viral titration

HEK293 cells were seeded in 24-well plates at a density of 2 × 10^5^ cells/mL and cultured for 18-24 h. Viral supernatants were serially diluted 10-fold, and each dilution was tested in triplicate. After 2 h of infection, the inoculum was replaced with complete DMEM containing 10% FBS, and cells were incubated for an additional 48 h. Cells were fixed with 500 μL of methanol per well for 30 min, washed three times with PBS, and incubated with anti-adenovirus type 5 (Ad5) primary antibody, diluted in 1% BSA, at room temperature for 2 h. After washing, HRP-conjugated secondary antibody was added and incubated for 1 h. Plates were washed three times with PBS. TMB substrate and H₂O₂ (1:200 dilution) were added (250 μL/well) and the wells were incubated in the dark for 20 min. After washing, brown staining spots representing viral plaques were visualized under a microscope and counted to calculate viral titers. All experiments involving oncolytic adenovirus were conducted in a BSL-2 biosafety laboratory.

### Western blot analysis

Cells were seeded in 12-well plates at 5 × 10^5^ cells/mL and cultured for 18-24 h, followed by viral infection and/or drug treatment as indicated. Cells were lysed in RIPA buffer containing PMSF for 30 min on ice. Lysates were centrifuged at 12,000 × g for 5 min, and supernatants were mixed with 5× SDS loading buffer and denatured at 100°C for 10 min. Proteins were separated by SDS-PAGE, transferred to membranes, and blocked for 2 h at room temperature. Membranes were incubated with primary antibodies overnight at 4°C, followed by HRP-conjugated secondary antibodies (1:5000) for 1 h at room temperature. Signals were detected using enhanced chemiluminescence (ECL). Adenovirus type 5 (Ad5; in-house rabbit serum) and β-actin as loading control.

### In vivo animal studies

All animal experiments were approved by the Institutional Animal Care and Use Committee. 5–6-week-old BALB/c and BALB/c-nu (nude) mice were housed at 22- 24°C with 40-70% humidity under pathogen-free conditions and provided food and water ad libitum. 4T1 cells (1 × 10^7^ cells/mL) and NCI-H1299 cells (2 × 10^7^ cells/mL) were subcutaneously injected (100 μL) into both flanks of mice. When tumor volumes reached approximately 100 mm³, mice were randomly assigned to groups (n = 10 per group). For therapeutic studies, two doses of OAd-2B6 were administered via intratumoral injection. Cyclophosphamide (CTX) was administered intraperitoneally 48 h after each viral injection for three consecutive doses. Tumor volume and body weight were measured every two days. Tumor size was calculated using the formula: V = (length × width^2^) / 2. Mice were euthanized when tumor volume exceeded 2000 mm^3^ or at endpoint. Tumor-free mice were defined as complete responders. Organs (heart, lung, liver, spleen, and kidney) and tumor tissues were collected for histological or molecular analyses.

### Immune profiling analysis

On days 7 and 14 post-treatment, tumors, spleens, axillary lymph nodes and inguinal lymph nodes were collected from 4T1 tumor-bearing mice (n = 3 per group). Single-cell suspensions from spleen and lymph nodes were prepared using lymphocyte separation medium. Tumor tissues were digested with collagenase IV (1 mg/mL), hyaluronidase (0.5 mg/mL), and DNase I (50 μg/mL) at 37°C for 1 h. Tumor-infiltrating lymphocytes were isolated using density gradient separation. Cells were stimulated with brefeldin A (1:1000, eBioscience^TM^) for 4 h at 37°C to block cytokine secretion. Dead cells were excluded using LIVE/DEAD^TM^ staining (Thermo Fisher Scientific), and Fc receptors were blocked using anti-mouse CD16/32 antibody (BD, 553142). Cells were fixed and permeabilized using BD fixation/permeabilization buffers (554714 and 554723). Fluorescent antibodies included CD45 (30-f11, 550994, BD) CD3e (SP34-2, 562877, BD), CD8α (53-6.7, 17-0081-82, eBioscience), CD4 (RM4-5, 552775), IFN-γ (XMG1.2, 505808, BioLegend), Granzyme B (NGZB, 11-8898-82, Invitrogen), CD25 (3C7, 564370, BD) and Foxp3 (NRRF-30, 14-4771-80, eBioscience). Samples were analyzed by flow cytometry with gating based on unstained and FMO controls.

### IFN-γ ELISPOT assay

ELISPOT plates were pre-wetted with 75% ethanol diluted in PBS (1:1), washed, and coated with anti-mouse IFN-γ capture antibody (1:200 in PBS) overnight at 4°C. After blocking with 5% R10 medium for 2 h, 3 × 10^6^/mL splenocytes were added and stimulated with tumor cell lysates for 20 h at 37°C in 5% CO_2_. Detection was performed using biotinylated anti-IFN-γ antibody followed by streptavidin-HRP/AP (1:2500). Spots were developed using AEC or BCIP/NBT substrate and counted using an ELISPOT reader. Specific spot counts were calculated by subtracting background (unstimulated wells).

### Cell viability assay

Cells were seeded in 96-well plates at a density of 2 × 10⁵ cells/mL and incubated for 18-24 h. Cells were then treated with OAd-2B6 and/or CTX for 48 or 72 h. CCK-8 reagent (10:1 with medium) was added, and the plate was incubated for 1 h. Absorbance was measured at 450 nm using a microplate reader.

### Statistical analysis

All data were analyzed using GraphPad Prism 9. Statistical significance was assessed using unpaired two-tailed t-test, one- or two-way ANOVA with Tukey’s post hoc test, Kaplan–Meier survival analysis (log-rank test), and Pearson correlation analysis, as appropriate. Data are presented as mean ± standard deviation (SD). Statistical significance was defined as: *P < 0.05, **P < 0.01, ***P < 0.001, ****P < 0.0001.

## Acknowledgements

This study was supported by the Project of Guangzhou Science and Technology (2025B01J3001) and Guangzhou nBiomed Co., Ltd.

## Author Contributions

L.C., L.Q., A.H. and S.G. conceived the idea, designed the experiments, and interpreted the data. M.S., H.Z., D.X., H.L., C.Y., S.G., and C.W. performed the experiments and collected the data. M.S., S.G., J.L., P.L., A.H., L.Q., and L.Q. prepared and revised the manuscript. All authors have read and approved the final manuscript.

## References

1. Raimondi V, Vescovini R, Dessena M, Donofrio G, Storti P, Giuliani N. Oncolytic viruses: a potential breakthrough immunotherapy for multiple myeloma patients. Front Immunol. 2024;15:1483806. 10.3389/fimmu.2024.1483806

2. Kelly E, Russell SJ. History of Oncolytic Viruses: Genesis to Genetic Engineering. Molecular Therapy. 2007;15:651–9. 10.1038/sj.mt.6300108

3. Andtbacka RHI, Collichio F, Harrington KJ, Middleton MR, Downey G, Ӧhrling K, et al. Final analyses of OPTiM: a randomized phase III trial of talimogene laherparepvec versus granulocyte-macrophage colony-stimulating factor in unresectable stage III–IV melanoma. j immunotherapy cancer. 2019;7:145. 10.1186/s40425-019-0623-z

4. Alberts P, Tilgase A, Rasa A, Bandere K, Venskus D. The advent of oncolytic virotherapy in oncology: The Rigvir® story. European Journal of Pharmacology. 2018;837:117–26. 10.1016/j.ejphar.2018.08.042

5. Liang M. Oncorine, the World First Oncolytic Virus Medicine and its Update in China. CCDT. 2018;18:171–6. 10.2174/1568009618666171129221503

6. Chen C, Cillis J, Deshpande S, Park AK, Valencia H, Kim SI, et al. Oncolytic Virotherapy in Solid Tumors: A Current Review. BioDrugs. 2025;39:857–76. 10.1007/s40259-025-00743-z

7. Du W, Na J, Zhong L, Zhang P. Advances in preclinical and clinical studies of oncolytic virus combination therapy. Front Oncol. 2025;15:1545542. 10.3389/fonc.2025.1545542

8. Kaufman HL, Kohlhapp FJ, Zloza A. Oncolytic viruses: a new class of immunotherapy drugs. Nat Rev Drug Discov. 2015;14:642–62. 10.1038/nrd4663

9. Ning W, Qian X, Dunmall LC, Liu F, Guo Y, Li S, et al. Non-secreting IL12 expressing oncolytic adenovirus Ad-TD-nsIL12 in recurrent high-grade glioma: a phase I trial. Nat Commun. 2024;15:9299. 10.1038/s41467-024-53041-7

10. Block MS. The oncolytic adenovirus TILT-123 with pembrolizumab in platinum resistant or refractory ovarian cancer: the phase 1a PROTA trial. 2025; 16:1381. 10.1038/s41467-025-56482-w

11. Uchio EM, Lamm DL, Shore ND, Kamat AM, Tyson M, Tran B, et al. A phase 3, single-arm study of CG0070 in subjects with nonmuscle invasive bladder cancer (NMIBC) unresponsive to Bacillus Calmette-Guerin (BCG). JCO. 2022;40:TPS598– TPS598. 10.1200/JCO.2022.40.6_suppl.TPS598

12. Yamanaka M, Tada Y, Kawamura K, Li Q, Okamoto S, Chai K, et al. E1B-55 kDa- Defective Adenoviruses Activate p53 in Mesothelioma and Enhance Cytotoxicity of Anticancer Agents. Journal of Thoracic Oncology. 2012;7:1850–7. 10.1097/JTO.0b013e3182725fa4

13. Xia ZJ, Chang JH, Zhang L, Jiang WQ, Guan ZZ, Liu JW, et al. Phase III randomized clinical trial of intratumoral injection of E1B gene-deleted adenovirus (H101) combined with cisplatin-based chemotherapy in treating squamous cell cancer of head and neck or esophagus. Ai Zheng. 2004; 23:1666–1670.

14. Li R, Steinberg G, Uchio E, Lamm D, Paras S, Kamat A, et al. 666 Phase 2, single arm study of CG0070 combined with pembrolizumab in patients with non-muscle invasive bladder cancer (NMIBC) unresponsive to bacillus calmette-guerin (BCG). Regular and Young Investigator Award Abstracts. BMJ Publishing Group Ltd; 2022. p. A696–A696. 10.1136/jitc-2022-SITC2022.0666

15. Makower D, Rozenblit A, Kaufman H, Edelman M, Lane ME, Zwiebel J, et al. Phase II Clinical Trial of Intralesional Administration of the Oncolytic Adenovirus ONYX-015 in Patients with Hepatobiliary Tumors with Correlative p53 Studies. Clin Cancer Res. 2003; 9: 693–702.

16. Zhu Z, McGray AJR, Jiang W, Lu B, Kalinski P, Guo ZS. Improving cancer immunotherapy by rationally combining oncolytic virus with modulators targeting key signaling pathways. Mol Cancer. 2022;21:196. 10.1186/s12943-022-01664-z

17. Voelcker G. Cyclophosphamide: Old Drug with Great Future. DDC. 2025;4:48. 10.3390/ddc4040048

18. Pol JG, Atherton MJ, Stephenson KB, Bridle BW, Workenhe ST, Kazdhan N, et al. Enhanced immunotherapeutic profile of oncolytic virus-based cancer vaccination using cyclophosphamide preconditioning. J Immunother Cancer. 2020;8:e000981. 10.1136/jitc-2020-000981

19. Chang KH, Weber GF, Crespi CL, Waxman DJ. Differential Activation of Cyclophosphamide and Ifosphamide by Cytochromes P-450 2B and 3A in Human Liver Microsomes. Cancer Res. 1993;53: 5629–5637.

20. Chen L, Waxman DJ. Cytochrome P450 Gene-directed Enzyme Prodrug Therapy (GDEPT) for Cancer. Curr Pharm Des. 2002;8: 1405–1416. 10.2174/1381612023394566

21. Braybrooke JP, Slade A, Deplanque G, Harrop R, Madhusudan S, Forster MD, et al. Phase I Study of MetXia-P450 Gene Therapy and Oral Cyclophosphamide for Patients with Advanced Breast Cancer or Melanoma. Clinical Cancer Research. 2005;11:1512–20. 10.1158/1078-0432.CCR-04-0155

22. Hu W, Bian Y, Ji H. TIL Therapy in Lung Cancer: Current Progress and Perspectives. Advanced Science. 2024;11:2409356. 10.1002/advs.202409356

23. Ji T, Li L, Li W, Zheng X, Ye X, Chen H, et al. Emergence and characterization of a putative novel human adenovirus recombinant HAdV-C104 causing pneumonia in Southern China. Virus Evolution. 2021;7:veab018. 10.1093/ve/veab018

24. Jiang H, Clise-Dwyer K, Ruisaard KE, Fan X, Tian W, Gumin J, et al. Delta-24- RGD Oncolytic Adenovirus Elicits Anti-Glioma Immunity in an Immunocompetent Mouse Model. Castro MG, editor. PLoS ONE. 2014;9:e97407. 10.1371/journal.pone.0097407

25. Fueyo J, Alemany R, Gomez-Manzano C, Fuller GN, Khan A, Conrad CA, et al. Preclinical Characterization of the Antiglioma Activity of a Tropism-Enhanced Adenovirus Targeted to the Retinoblastoma Pathway. JNCI Journal of the National Cancer Institute. 2003;95:652–60. 10.1093/jnci/95.9.652

26. Han J, Sabbatini P, Perez D, Rao L, Modha D, White E. The E1B 19K protein blocks apoptosis by interacting with and inhibiting the p53-inducible and death-promoting Bax protein. Genes Dev. 1996;10:461–77. 10.1101/gad.10.4.461

27. Pantelidou C, Cherubini G, Lemoine NR, Halldén G. The E1B19K-deleted oncolytic adenovirus mutant AdΔ19K sensitizes pancreatic cancer cells to drug-induced DNA-damage by down-regulating Claspin and Mre1. Oncotarget. 2016; 7:15703– 15724. 10.18632/oncotarget.7310.

28. Piya S, White EJ, Klein SR, Jiang H, McDonnell TJ, Gomez-Manzano C, et al. The E1B19K Oncoprotein Complexes with Beclin 1 to Regulate Autophagy in Adenovirus-Infected Cells. Fimia GM, editor. PLoS ONE. 2011;6:e29467. 10.1371/journal.pone.0029467

29. Lin D, Shen Y, Liang T. Oncolytic virotherapy: basic principles, recent advances and future directions. Sig Transduct Target Ther. 2023;8:156. 10.1038/s41392-023-01407-6

30. Wang W, Chu Y, Zhao L, Lv M, Wang L, Jin F, et al. HBsAg-tagged tumour vaccine system eliminates solid tumours through virus-specific memory T cells. Nat Biomed Eng. 2025 . 10.1038/s41551-025-01555-w

31. Kang X, Han Y, Wu M, Li Y, Qian P, Xu C, et al. In situ blockade of TNF-TNFR2 axis via oncolytic adenovirus improves antitumor efficacy in solid tumors. Molecular Therapy. 2025;33:670–87. 10.1016/j.ymthe.2024.12.011

32. Zhao Y, Liu Z, Li L, Wu J, Zhang H, Zhang H, et al. Oncolytic Adenovirus: Prospects for Cancer Immunotherapy. Front Microbiol. 2021;12:707290. 10.3389/fmicb.2021.707290

33. Sutherland TE, Dyer DP, Allen JE. The extracellular matrix and the immune system: A mutually dependent relationship. Science. 2023;379:eabp8964. 10.1126/science.abp8964

34. Ramesh N, Ge Y, Ennist DL, Zhu M, Mina M, Ganesh S, et al. CG0070, a Conditionally Replicating Granulocyte Macrophage Colony-Stimulating Factor– Armed Oncolytic Adenovirus for the Treatment of Bladder Cancer. Clinical Cancer Research. 2006;12:305–13. 10.1158/1078-0432.CCR-05-1059

35. Ayala G, Satoh T, Li R, Shalev M, Gdor Y, Aguilar-Cordova E, et al. Biological Response Determinants in HSV-tk + Ganciclovir Gene Therapy for Prostate Cancer. Molecular Therapy. 2006;13:716–28. 10.1016/j.ymthe.2005.11.022

